# Affinity purification-mass spectrometry and single fiber physiology/proteomics reveals mechanistic insights of C18ORF25

**DOI:** 10.1101/2023.10.30.564812

**Authors:** Yaan-Kit Ng, Ronnie Blazev, James W. McNamara, Mriga Dutt, Jeffrey Molendijk, Enzo R. Porrello, David A. Elliott, Benjamin L. Parker

**Affiliations:** Department of Anatomy & Physiology, The University of Melbourne, Parkville, VIC, Australia; Centre for Muscle Research, The University of Melbourne, Parkville, VIC, Australia; Murdoch Children’s Research Institute and Melbourne Centre for Cardiovascular Genomics and Regenerative Medicine, The Royal Children’s Hospital, Parkville, VIC, Australia; Melbourne Centre for Cardiovascular Genomics and Regenerative Medicine, The Royal Children’s Hospital, Melbourne 3052, VIC, Australia; Novo Nordisk Foundation Center for Stem Cell Medicine, Murdoch Children’s Research Institute, Melbourne 3052, VIC, Australia; Department of Paediatrics, Faculty of Medicine, Dentistry & Health Sciences, The University of Melbourne, Melbourne 3010, VIC, Australia

## Abstract

C18ORF25 was recently shown to be phosphorylated at S67 by AMP-activated protein kinase (AMPK) in skeletal muscle following acute exercise in humans. Phosphorylation was shown to improve *ex vivo* skeletal muscle contractile function in mice but our understanding of the molecular mechanisms is incomplete. Here, we profiled the interactome of C18ORF25 in mouse myotubes using affinity purification coupled to mass spectrometry. This analysis included an investigation of AMPK-dependent and S67-dependent protein:protein interactions. Several nucleocytoplasmic and contractile-associated proteins were identified, and revealed a subset of GTPases that associate with C18ORF25 in an AMPK- and S67 phosphorylation-dependent manner. We confirmed that C18ORF25 is localised to the nucleus and the contractile apparatus in skeletal muscle. Mice lacking *C18Orf25* display defects in calcium handling specifically in fast-twitch muscle fibers. To investigate these mechanisms, we developed an integrated single fiber physiology and single fiber proteomic platform. The approach enabled detailed assessment of various steps in the excitation-contraction (EC) pathway including SR calcium handling and force generation followed by paired single fiber proteomic analysis. This enabled us to identify >700 protein:phenotype associations and 36 fiber-type specific differences following loss of *C18Orf25*. Taken together, our data provide unique insights into the function of C18ORF25 and its role in skeletal muscle physiology.

## Introduction

The beneficial outcomes of exercise are achieved through a series of protein signalling networks ^1^. This is largely mediated via protein phosphorylation which involves the reversible, kinase catalysed, attachment of a phosphoryl group to a target protein which can then regulate its function. For example, depletion of ATP stores during exercise leads to the activation of AMP-activated protein kinase (AMPK). Multiple downstream substrates of AMPK are then phosphorylated which act to inhibit anabolic cellular processes while promoting catalytic reactions that result in the generation of ATP. Over time, repeated activation of AMPK induces gene transcription via several proteins that result in improvements in oxidative capacity and therefore metabolic profile^2^.

Over the past decade, several phosphoproteomic analyses of exercise in human and rodent models have revealed the regulation of thousands of phosphorylation sites ^3, 4, 5, 6, 7, 8, 9, 10, 11^. We previously performed a phosphoproteomic analysis of human skeletal muscle biopsies taken before, immediately after and three hours into recovery from different modalities of sprint, resistance, and endurance exercise ^11^. This enabled us to discover the canonical core set of exercise-regulated phosphorylation sites which likely underpin the benefits of exercise. Phosphorylation of S67 on the uncharacterized protein C18ORF25 was identified in this canonical signalling group and selected for further functional characterization.

C18ORF25 (also known as ARKadia (RNF111) N-terminal like PKA signalling regulator protein 2N [ARK2N]) is a homolog of Arkadia (RNF111), an E3 ubiquitin ligase with SUMO-interaction motifs (SIMs) ^12^. However, C18ORF25 lacks the entire C-terminal RING domain of RNF111 which is required for ubiquitin binding suggesting it lacks ubiquitination activity and may therefore act as an adaptor or signalling scaffold ^13^. We have previously shown that mice lacking *C18Orf25* throughout the entire body have increased adiposity, decreased lean mass, lower exercise capacity and significantly reduced *ex vivo* skeletal muscle force production ^11^. Skeletal muscle isolated from *C18Orf25* knockout (KO) mice have reduced cAMP-dependent protein kinase A (PKA) levels, and reduced phosphorylation of several contractile proteins and proteins involved in calcium handling. Furthermore, analysis of single muscle fibers from *C18Orf25* KO mice revealed impaired SR calcium cycling in fast-twitch fibers only ^11^. Critically, the exercise-regulated phosphorylation of S67 on C18ORF25 is directly catalysed by AMPK. This phosphorylation site plays an important role in the function of C18ORF25 as re-expression of a phospho-mimetic S66/67D but not phospho-dead S66/67A in *C18Orf25* KO mice is able to rescue skeletal muscle contractile defects ^11^.

In the present study, we further investigated the potential molecular functions of C18ORF25. We first performed affinity purification coupled to tandem mass spectrometry (AP-MS) to probe the protein:protein interaction (PPI) landscape of C18ORF25. Included in this analysis was also an investigation of potential AMPK- and S67 phosphorylation-dependent PPIs. These data promoted us to investigate the subcellular location of C18ORF25 and its role in *ex vivo* contraction-stimulated skeletal muscle glucose uptake. Our previous phenotypic analysis of single muscle fibers isolated from wild-type (WT) or *C18Orf25* KO mice highlighted defects in SR calcium handling specifically in fast-twitch (FT) fibers. Hence, we investigated these mechanisms by developing an integrated single fiber physiology and single fiber proteomic platform to identify hundreds of novel phenotype:protein correlations. The analysis also enabled us to identify proteome differences specifically in FT fibers following loss of C18ORF25. Taken together, our data suggest C18ORF25 is likely a multi-functional protein with several underlying mechanisms contributing to skeletal muscle physiology.

## Methods

### Mouse housing

All mouse experiments were approved by The University of Melbourne Animal Ethics Committee (AEC ID1914940) and conformed to the National Health and Medical Research Council of Australia guidelines regarding the care and use of experimental animals. *C18Orf25* WT and KO mice were generated and validated as described previously ^11^. Mice were housed at 22°C (+/−1°C) in groups of five/cage and maintained on a Standard Chow diet (Specialty Feeds, Australia) with a 12-h light/dark cycle and ad libitum access to food and water.

### Cell Culture and affinity purification sample preparation

C2C12 cells were grown in Dulbecco’s Modified Eagle Medium (DMEM) (GIBCO by Life Technologies; # 11995065), supplemented with 10% fetal bovine serum (FBS) (Life Technologies; #26140079), pyruvate and GlutaMAX (GIBCO by Life Technologies). Cells were kept at 37°C and 5% CO2 in a humidified incubator Direct Heat CO2 Incubator featuring Oxygen Control (In Vitro Technologies). Mouse C18Orf25 cDNA (8030462N17Rik; NM_178670.4) was purchased from VectorBuilder with an N-terminal 3xFLAG tag as either wild-type (WT) or S66/67A mutations and sub-cloned into a pAAV expression vector containing CMV promoter. The vectors sequences and mutations were confirmed by Sanger sequencing and restriction enzyme digestions ^11^. Vectors were transiently transfected at 60% confluence in a 6-well plate with 3 µg DNA, 9 µl of P3000 reagent and 9 µl of Lipofectamine 3000 prepared in reduced-serum Minimal Essential Medium (Opti-MEM) (Life Technologies). After 24 hours, the cells were differentiated into myotubes with the replacement of 10% FBS with 2% Horse Serum and cultured for a further 3 days. On the day of harvest, cells were incubated in 1ml of DMEM, Pyruvate, GlutaMAX with 0.2% BSA for 2 hours to achieve a basal state. Relevant cells were then stimulated for 30 minutes with 2mM AICAR in basal media and 100uM of A-769662 in DMSO. Cells were washed twice with cold PBS, then lysed in 200ul of interactome lysis buffer (0.3% CHAPS, 150mM NaCl, 50mM Tris pH 7.5, 5% glycerol, EDTA-Free PIC, PhosIC). Lysates were pushed through a 22G needle 6 times followed by a 27G needle 3 times. Lysates were spun down at 18,000 g for 10 minutes and normalised to 700 µg of protein as determined with BCA (ThermoFisher Scientific). 1.5ul of mouse anti-FLAG M2 (Sigma #F1804) was added to each lysate and incubated for 2 hours with rotation at 4°C. 40µl of protein-G beads (Invitrogen; #10004D) was washed with wash buffer II (150mM NaCl, 50mM Tris pH 7.5, 5% glycerol) on the DynaMag-2 magnet (ThermoFisher Scientific) and added to each lysate and rotated for 2 hours at 4°C. The beads were then washed twice with 950ul of wash buffer I (0.01% CHAPS in wash buffer II), followed by two washes in 1ml of wash buffer II leaving the final wash solution with the beads. 50ul of beads was then removed for western blot where proteins were eluted with 50ul of 2x Laemmli + TCEP buffer. The remaining final wash solution was then removed after which we performed an on-bead digestion by adding 50ul digestion buffer (2M urea, 50mM Tris pH 7.5 containing 1 mM TCEP, 5mM CAA) as well as 0.2ug of Trypsin (Promega) and LysC (Wako) overnight at 37°C. Samples were then acidified to a final concentration of 1% trifluoroacetic acid (TFA) in 90% isopropanol and purified by in-house packed SDB-RPS (Sigma; #66886-U) microcolumns. The purified peptides were resuspended in 2% acetonitrile in 0.1% TFA and stored at −80°C prior to direct injection by LC-MS/MS.

### *Ex vivo* Contraction-Stimulated Glucose Uptake

Mice were anesthetized with either pentobarbitone sodium (270 mg/kg body weight via intraperitoneal injection) and transferred to a dissecting stage. Depth of anesthesia was assessed via the lack of leg and optical reflexes. After confirming anesthesia, skin from the hind legs was removed, and *extensor digitorum longus* (EDL) muscles were sutured using 5.0 braided suture at both proximal and distal ends at the tendomuscular junction. Muscles were excised and incubated in Modified Krebs Buffer (MKB) (116 mM NaCl, 4.6 mM KCl, 1.16 mM KH2PO4, 25.3 mM NaHCO3, 2.5 mM CaCl2, 1.16 mM MgSO4) in a myograph (DMT, Denmark; #820MS) at 30°C with constant gentle bubbling of 5% medical carbon dioxide in oxygen.

Left and right EDL muscles were adjusted to resting tension (∼5 mN) and allowed to equilibrate for 10 min before being electrically contracted (left leg basal, right leg stimulated) for 10 min. Here, a DMT CS4 stimulator was used to deliver 0.2 ms supramaximal (26V) pulses via stimulation electrodes (DMT 300145) placed over the mid-belly of the muscle. Right muscles were stimulated at 60 Hz, 350 ms duration, 1 every 5 s for 10 min. Muscles were left for another 10 min after which the buffer was replaced with MKB containing 0.375 μCi/ml 2-deoxy-d-[1,2-3H] glucose and 0.05 μCi/ml d-[14C] mannitol for another 10 min. Muscles were then immediately immersed in ice-cold PBS to stop further glucose uptake. Each muscle was then excised of its sutures/tendons and frozen in liquid nitrogen for future lysis in 400 μl of 2% SDS via tip-probe sonication (QSonica). Measurement of radiolabelled 2-deoxyglucose was carried out in a Tri-Carb 4910 TR liquid scintillation analyzer (Perkin Elmer) by adding 200 μl of muscle lysate to 4 ml scintillation fluid (Ultima Gold, Perkin Elmer). Glucose uptake rates were calculated as described in ^14^.

### Western Blotting

Protein was separated on NuPAGE 4-12% Bis-Tris protein gels (ThermoFisher Scientific) in MOPS SDS Running Buffer at 160 V for 1 h at room temperature. The protein was transferred to PVDF membranes (Millipore; #IPFL00010) in NuPAGE Transfer Buffer at 20 V for 1 h at room temperature and blocked with 5% skim milk in Tris-buffered saline containing 0.1% Tween 20 (TBST) for at least 30 min at room temperature with gentle shaking. The membranes were incubated overnight in primary antibody with 5% BSA in TBST with gentle roller at 4°C and washed three times in TBST at room temperature. The membranes were incubated with HRP-secondary antibody in 5% skim milk in TBST for 45 min at room temperature and washed three times with TBST. Protein was visualized with Immobilon Western Chemiluminescent HRP Substrate (Millipore; #WBKLS0500) and imaged on a ChemiDoc (BioRad). Densitometry was performed in ImageJ ^15^.

### Immunofluorescence microscopy

Tibialis anterior (TA) muscles from WT mice were injected with either AAV6-C18ORF25 or AAV6-empty vector to assess localization of C18ORF25 in striated muscles. After euthanasia, TA muscles were excised and cryopreserved in OCT. 5μm thick longitudinal sections were made using a cryostat at approximately the mid-belly of the TA and slides stored at -80°C. For staining of sections, slides were warmed to room temperature prior to fixation with 4% paraformaldehyde in PBS for 10 minutes. Sections were then blocked and permeabilized in 5% goat serum, 2% BSA, 0.1% triton X-100 in PBS. Sections were then stained overnight at 4°C with 1:1000 anti-alpha actinin (Abcam ab68167) and 1:250 anti-FLAG (Sigma F1804) in PBS containing 0.05% Tween-20. After washing, primary antibodies were labelled with goat anti-mouse IgG Alexa Fluor 555 (A21422) and goat anti-rabbit IgG Alexa Fluor 488 (A11034) and counterstained with DAPI. Images were collected at 20x on a Zeiss LSM900.

### Single fiber physiology

Mice were anesthetized with pentobarbitone sodium (270 mg/kg body weight) and transferred to a dissecting microscope stage. Depth of anaesthesia was assessed via the lack of leg and optical reflexes. After confirming anaesthesia, skin from the hind legs was removed, and *soleus* muscles were quickly harvested and pinned at resting length in Paraffin oil in a Petri dish lined with Sylgard 184. Single mechanically skinned muscle fibers were prepared and utilized as described in Lamb and Stephenson ^16^. Briefly, under a dissecting microscope (Olympus SZ61) and with the aid of fine jeweller forceps and micro-scissors (Vannas), a segment of a single muscle fiber was dissected free and then mechanically-skinned by removing its surface membrane. A video camera (Panasonic WV-CP474) monitor system attached to the microscope was used to calculate the cross-sectional area (CSA) of the skinned segment (assumed to be cylindrical) according to the formula CSA (μm2) = π(d/2 x a)2 where d is the diameter of the skinned segment and a is the magnification conversion factor for diameter. The skinned fiber was then attached to a force transducer (AME801, Norway) at one end using 10-0 suture silk (Genzyme) and clamped with fine forceps at the other end, before being stretched to 120% of its resting length and immersed in a 2 mL bath containing the standard K-HDTA solution (mimicking the intracellular milieu of an intact fiber). Force responses were digitized using a PowerLab 2/25 unit (ADInstruments, Australia) and analyzed with the Peak Parameters module in LabChart Pro (v8.1.16, ADInstruments). On the day of experiments, the Petri dish containing the muscle was kept at 5°C when not in use and all experiments were performed at 20 ± 1°C.

The standard K-HDTA solution (referred hereon as 3NF) comprised in mM: 126 K+, 36 Na+, 8.5 total Mg2+ to give 1 free [Mg2+], 90 HEPES, 50 HDTA (1,6-Diaminohexane-N,N,N′,N′-tetraacetic acid), 0.025 total EGTA, 8 ATP, 10 CP, with pH 7.10. The INF (relaxing, pCa (-log10[Ca2+] >9) and IINF (maximum Ca2+-activating, pCa 4.48) solutions used for contractile apparatus experiments were similar to the standard K-HDTA solution except all HDTA was replaced with 50 mM EGTA (INF) or 50 mM Ca-EGTA (IINF) and total Mg2+ changed to 10.3 and 8.1 mM, respectively, to give 1 mM free [Mg2+]. These INF and IINF solutions could be mixed in specific ratios to give the desired free [Ca2+] required for the Ca2+-sensitivity experiments described below. A strontium solution with pSr (-log10[Sr2+]) 5.3 was used to differentiate between type I (slow-twitch) and type II (fast-twitch) fibers ^17^.

To assess SR Ca^2+^ handling, the skinned fiber was exposed for 10 s to 3NF with 0.5 mM EGTA before the SR was fully depleted of all its releasable Ca2+ content by exposure for 1 min to 3NF with 30 mM caffeine, 0.05 mM free Mg2+ and 0.5 mM EGTA (i.e. release solution). The time-integral (area) of the resulting caffeine-induced force response was indicative of the amount of releasable Ca2+ present in the SR immediately prior to caffeine exposure ^16^, and thus the endogenous SR Ca2+ content given fibers were always skinned under paraffin oil and therefore retained their endogenous SR Ca2+ levels. Following this, the skinned fiber was washed for 1 min in 3NF with 1 mM EGTA to remove caffeine from the preparation and prevent any SR Ca2+ reuptake. The SR could then be reloaded with Ca2+ to a set level by exposure to a Ca2+ load solution (3NF with 1 mM total EGTA-CaEGTA, pCa 7.0) for a set period (15–120 s), before again being depleted by exposure to the (30 mM caffeine) release solution. Such load-deplete cycles could be repeated with the area under 30 mM caffeine force response used as an indicator of the amount of Ca2+ that could be reloaded into the SR. To estimate the extent of Ca2+ loss from the SR due to passive leak, the SR was first depleted of Ca2+ as described above and then reloaded with Ca2+ for 1 min before being depleted again. The SR was then loaded for 1 min with Ca2+, incubated for 1 min in a leak solution (3NF with 1 mM EGTA to chelate any Ca2+ passively leaking from the SR and prevent Ca2+ reuptake by the SR Ca2+ pump) and subsequently depleted. The area of this last force response gave an indication of the amount of Ca2+ present in the SR immediately after the 1 min leak period and could be compared to the area of the previous response with no leak (i.e. area of post-leak vs area of pre-leak response).

Upon completion of the SR Ca2+ experiments, contractile function was assessed by exposing the skinned fiber to INF with 2% Triton-X100 for 5 min to destroy all membranous compartments and then washed twice for 2.5 min in INF without Triton. The fiber was subsequently exposed to the pSr 5.3 solution for fiber type determination, before being relaxed in INF and then exposed to IINF to ascertain the maximum Ca2+-activated force response (Fmax) the contractile apparatus could generate. The Ca^2+^-sensitivity of the contractile apparatus was assessed next by exposing the skinned fiber to a sequence of solutions (mixtures of INF and IINF) in which the free [Ca2+] was progressively increased to higher levels (pCa 7.0 to 4.48) until Fmax was reached, after which the fiber was relaxed completely by exposure to INF. The force produced at each Ca2+ concentration for the Force-pCa staircase just described could be normalized to Fmax and plotted as a function of pCa using GraphPad Prism. From Hill curves fitted to plots of force versus pCa for each fiber, it was possible to obtain the parameters pCa50 (pCa giving 50% Fmax, an indicator of the Ca2+-sensitivity of the contractile apparatus) and Hill coefficient (an indicator of the steepness of the Force-pCa curve).

Upon completion of single fibre physiology experiments, individual fibers were collected in 50 µl of 2% sodium dodecyl sulfate in 100 mM Tris pH 8.5 and snap frozen.

### Single fiber proteomic sample preparation

Single fibers were reduced with 10 mM tris(2-carboxyethyl)phosphine (Sigma; #75259) and 40 mM 2-chloroacetamide (Sigma; #22790) in 100 µl of 1% SDS in 100 mM Tris, pH 8.5 and heated at 95°C for 5 min then lysed via tip-probe sonication. Samples were prepared using a modified SP3 protocol ^18^. Briefly, Seramag speedbead carboxyl beads (GE Life Sciences) were prepared where a 1:1 mix of hydrophilic (Cat No. 45152105050250) and hydrophobic (Cat No. 65152105050250) beads were washed with and resuspended in LC grade H2O at 10mg/ml of each bead type. 10ul of prepared beads and 110ul of LC grade ethanol were then added to each lysate. They were then shaken for 8 minutes at 1600RPM, spun down at 1000xg for 1 minute at room temp. Lysates were placed on a DynaMag-2 magnet (ThermoFisher Scientific) for 1 minute after which supernatant was discarded. Beads were then washed 3x with 950ul of 80% LC grade ethanol. The last wash was removed, and beads were air dried for approximately 10 minutes then resuspended in 25ul of Digestion Buffer (10% 2,2,2-Trifluoroethanol (Sigma; #96924)) in 100 mM HEPEs pH 8.5). Lysates were digested with 60 ng of sequencing grade trypsin (Sigma; #T6567) and 60 ng of sequencing grade LysC (Wako; #129-02541) overnight at 37°C with shaking at 2000 RPM. Samples were then acidified to a final concentration of 1% TFA in 90% isopropanol and purified by in-house packed SDB-RPS (Sigma; #66886-U) microcolumns. The purified peptides were resuspended in 2% acetonitrile in 0.1% TFA and stored at −80°C prior to direct injection by LC-MS/MS.

### LC-MS/MS acquisition

For analysis of AP-MS, peptides were analyzed on a Dionex 3500 nanoHPLC, coupled to an Orbitrap Exploris 480 mass spectrometer (ThermoFischer Scientific) via electrospray ionization in positive mode with 2.1 kV at 275 °C and RF set to 40%. Separation was achieved on a 50 cm × 75 µm column packed with C18AQ (1.9 µm; Dr Maisch, Ammerbuch, Germany) (PepSep, Marslev, Denmark) over 60min at a flow rate of 300 nL/min. The peptides were eluted over a linear gradient of 3–40% Buffer B (Buffer A: 0.1% v/v formic acid; Buffer B: 80% v/v acetonitrile, 0.1% v/v FA) and the column was maintained at 50 °C. The instrument was operated in data-independent acquisition mode with an MS1 spectrum acquired over the mass range 350–951 m/z (60,000 resolution, 250% automatic gain control (AGC) and 50 ms maximum injection time) followed by MS/MS via HCD fragmentation with 37 windows with 16 *m/z* isolation and a 1 *m/z* overlap (30,000 resolution, 2000% AGC and maximum injection time set to auto). For analysis of single fibers, peptides were analyzed on a Dionex 3500 nanoHPLC, coupled to an Orbitrap Exploris 480 mass spectrometer (ThermoFischer Scientific) via electrospray ionization in positive mode with 1.9 kV at 275 °C and RF set to 40%. Separation was achieved on a 50 cm × 75 µm column packed with C18AQ (1.9 µm; Dr Maisch, Ammerbuch, Germany) (PepSep, Marslev, Denmark) over 90min at a flow rate of 300 nL/min. The peptides were eluted over a linear gradient of 3–40% Buffer B (Buffer A: 0.1% v/v formic acid; Buffer B: 80% v/v acetonitrile, 0.1% v/v FA) and the column was maintained at 50 °C. The instrument was operated in data-independent acquisition mode with an MS1 spectrum acquired over the mass range 350–1400 m/z (60,000 resolution, 250% automatic gain control (AGC) and 50 ms maximum injection time) followed by MS/MS via HCD fragmentation with 50 windows with 13.7 *m/z* isolation and a 1 *m/z* overlap (30,000 resolution, 2000% AGC and maximum injection time set to auto).

### LC-MS/MS Data Processing and analysis

AP-MS were searched against the UniProt mouse database (October 2020; UP000000589_109090 and UP000000589_109090_additional) with Spectronaut v15.0.210615.50606 using library-free directDIA with default parameters and peptide spectral matches, peptide and protein false discovery rate (FDR) set to 1%. All data were searched with oxidation of methionine and N-terminal protein acetylation set as the variable modification and carbamidomethylation set as the fixed modification. Peptide quantification was carried out at MS2 level using 3-6 fragment ions, with automatic interference fragment ion removal as previously described ^19^. The MS1 mass tolerance was set to 20 ppm, while the mass tolerance for MS/MS fragments was set to 0.02 Da. Dynamic mass MS1 and MS2 mass tolerance was enabled, and retention time calibration was accomplished using local (non-linear) regression. A dynamic extracted ion chromatogram window size was performed. Analysis of single fiber proteomics data was performed with identical settings described above except against UniProt mouse database released in May 2022 and with Spectronaut v16.0.220606.53000. Data were analysed in Perseus ^20^ with Log_2_(x) transformation and normalisation by median subtraction. Differential abundance was calculated with Student’s t-test and q-values generated using Benjamini-Hochberg with FDR set to 5%. Pathway enrichment analysis was performed with Metascape ^21^. Protein:phenotype correlations were performed in CoffeeProt ^22^. Briefly, proteins were filtered to quantification in at least 50% of fibers (958 out of 1,612 proteins) and then pair-wise correlation performed to the phenotypes using biweight midcorrelation (bicor) as implemented in the WGCNA R package and corrected for multiple hypothesis testing using Benjamini-Hochberg ^23^.

## Data availability

Raw proteomic data and Spectronaut search results are available on the ProteomeXchange via the PRIDE database. The AP-MS data are available with the ProteomeXchange accession: PXD045629 (Reviewer username; password: pCkWUO04). The single fiber proteomic data are available with the ProteomeXchange accession: PXD045631 (Reviewer username; password: OdhYKkh3).

## Acknowledgements

We thank Nicholas Williamson, Ching-Seng Ang, Shuai Nie, Swati Varshney and Michael Leeming for instrument support in the Bio21 Mass Spectrometry and Proteomics Facility. This research was supported by access to the Melbourne Mouse Metabolic Phenotyping Platform at The University of Melbourne. This work was funded by an NHMRC Emerging Leader Investigator Grant (APP2009642), a Diabetes Australia Grant, a University of Melbourne Driving Research Momentum Grant, and an Australian Research Council (ARC) Discovery Project (DP230102652) to B.L.P. This work was also supported by a Department of Anatomy and Physiology (The University of Melbourne) ECR Seeding Grant to R.B. Y.-K.N. is a recipient of a School of Biomedical Science Postgraduate Award. The generation of the *C18Orf25* knockout mice used in this study was supported by the Phenomics Australia (PA) and the Australian Government through the National Collaborative Research Infrastructure Strategy (NCRIS) program. E.R.P. is supported by an Investigator Grant from the National Health and Medical Research Council (GNT 2008376). The Novo Nordisk Foundation Center for Stem Cell Medicine (E.R.P., D.A.E) is supported by Novo Nordisk Foundation grants (NNF21CC0073729). The Murdoch Children’s Research Institute is supported by the Victorian Government’s Operational Infrastructure Support Program.

## Results & Discussion

### The C18ORF25 interactome and S67 phosphorylation-dependent protein:protein interactions (PPIs)

To identify interacting partners of C18ORF25 and those regulated by AMPK-dependent phosphorylation of S67, we performed anti-FLAG affinity purification coupled to mass spectrometry (AP-MS) of differentiated C2C12 myotubes expressing either GFP, FLAG-C18ORF25-WT or FLAG-C18ORF25-S66/67A that were either untreated as controls (C) or dual treated with the AMPK activators AICAR and A-769662 (AA) (**Figure 1A**). Western-blot analysis confirmed AA-stimulated phosphorylation of S79 on Acetyl-CoA Carboxylase (ACC) and S67 on C18ORF25, with immunoreactivity lost following mutation of S66/67A (**Figure 1B**). AP-MS quantified 2,456 proteins with 380 significantly enriched with WT-Control vs GFP and/or WT-AA vs GFP (q<0.05 with Benjamini Hochberg FDR) (**Table S1**). **Figure 1C** plots these 380 proteins expressed as Log2(WT-Control/GFP) vs Log2(WT-AA/GFP) highlighting successful enrichment of C18ORF25 with the highest fold-change. Top interacting proteins include several Casein Kinase 2 isoforms and nuclear proteins such as transcription facotrs FOXK2, JUN and JUND, and the splicing factor PRPF31. Other top interacting proteins include the lipid kinase PIP4K2B, the translation initiation factor EIF3J1/2, and the uncharacterised proteins GPATCH2 (C14ORF118) and FAM76B. Of the 380 enriched proteins, 290 have been identified as SUMOylated suggesting the SUMO-interaction motif of C18ORF25 play an important role in protein:protein interactions ^24^. Functional annotation revealed an enrichment of proteins associated with endocytosis, regulation of the actin cytoskeleton, ribosomal proteins, mitophagy, lipid and atherosclerosis proteins, ferroptosis, adherens junctions, dopaminergic synapse as well as RAS, RAP1, Relaxin and Estrogen signalling pathways (**Figure 1D**). These data suggest possible wide-spread functions of C18ORF25.

**Figure 1.**
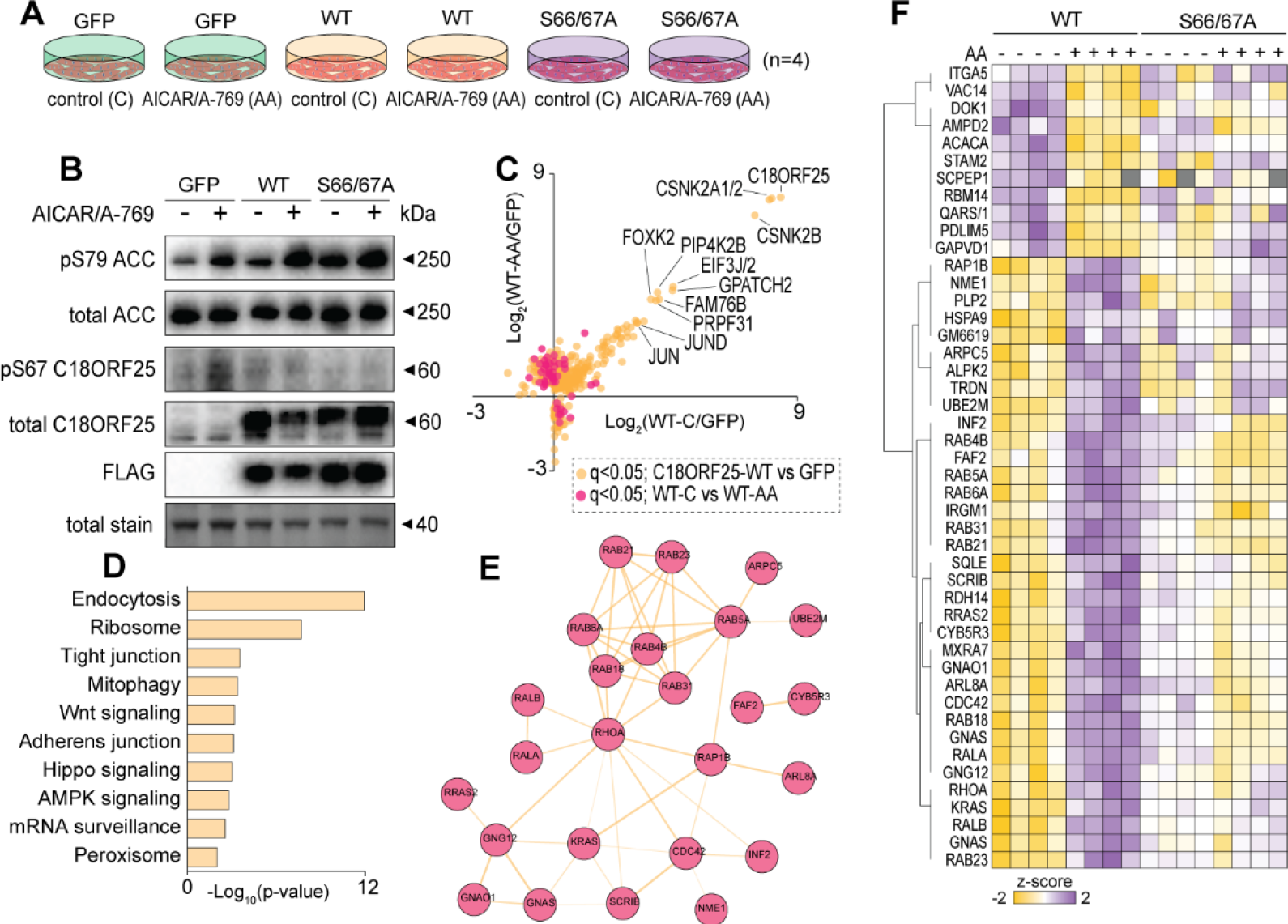
Affinity purification coupled to tandem mass spectrometry to profile the interactome, and AMPK- and S67 phosphorylation-dependent protein interactions of murine C18ORF25 in C2C12 myotubes. (**A**) Overview of experimental design. (**B**) Western-blot analysis of cells expressing i) GFP negative control, ii) N-FLAG-C18ORF25 wild-type (WT), or iii) N-FLAG-C18ORF25 S66/67A mutant stimulated with vehicle control (C) or with the AMPK activators AICAR/A-769662 (AA). (**C**) Scatterplot of enriched proteins in cells transfected with WT relative to GFP negative control cells. (**D**) Pathway analysis of enriched proteins associated with N-FLAG-C18ORF25-WT that display differential association following AMPK activation with AICAR/A-769662. (**E**) String-DB analysis of proteins displaying differential enrichment to N-FLAG-C18ORF25-WT following AMPK activation with AICAR/A-769662. (**F**) Heatmap of proteins displaying differential expression in N-FLAG-C18ORF25-S66/67A relative to WT.

Next, we investigated proteins that display differential enrichment to wild-type C18ORF25 in response to AMPK activation. Of the 380 interacting proteins, a total of 46 proteins were differentially enriched following AA stimulation which are indicated as red dots in **Figure 1C** and shift away from the diagonal on the scatterplot. The majority of these proteins increase their association with C18ORF25 following AMPK activation and shift upwards on the vertical axis. Interestingly, these proteins all display comparatively low enrichment suggesting sub-stoichiometric interactions as none of the top enriched proteins appear to differentially bind C18ORF25 in response to AMPK activation. Analysis of protein:protein associations in the STRINGdb revealed an interconnected network of GTPases (**Figure 1E**). This includes Ras (KRAS and RRAS2), Ras-related proteins such as several Rabs, and other multifunctional GTPases such as RALA/B, RHOA and CDC42. Additional proteins in this network include guanine nucleotide-binding proteins such as GNAS and GNAO1, and other proteins known to regulate GTPase activity such as SCRIB.

We next investigated S67 phosphorylation-dependent PPIs. Of the 46 proteins interacting with C18ORF25 in an AMPK-dependent manner, 26 display differential enrichment in the S66/67A mutant (**Figure 1F**). The majority of these proteins show increased association with wild-type C18ORF25 following stimulation of AMPK with AA but fail to increase association in the C18ORF25-S66/67A mutant suggesting their interactions require S67 phosphorylation. These proteins were primarily within the interconnected network of GTPases and their regulators identified from the STRINGdb. Additional proteins not directly within this network were also identified as AMPK- and S67 phosphorylation-dependent interactors of C18ORF25 such as SQLE, a squalene monooxygenase. SQLE is the rate-limiting enzyme in the synthesis of cholesterol in the mevalonate pathway. The mevalonate pathway also produces the precursors geranylgeranyl pyrophosphate and farsenyl pyrophosphate required for protein prenylation ^25^. Furthermore, SQLE can also directly generate farsenyl pyrophosphate from lanosterol. This is interesting because >15 proteins identified as AMPK- and S67 phosphorylation-dependent interactors of C18ORF25 are annotated as prenylated in the UniProt database including the previously mentioned network of GTPases where prenylation states are known to regulate important metabolic processes such as GLUT4 translocation ^26^. PDLIM5, otherwise known as ENH1, of the enigma family of proteins is another interesting interactor primarily characterised in cardiomyocytes. It is highly expressed in skeletal muscle, known to co-localize with α-actinin and is required for healthy myogenesis ^27^. PDLIM5 has also been shown to scaffold PKC/PKD1 with CREB via its PDK and LIM domains to facilitate phosphorylation of CREB at Ser133 and therefore CREB gene transcription ^28^. Furthermore, PDLIM5 is known to form various protein complexes at the z-disk to maintain sarcomeric structural integrity where they may interact with the extracellular matrix (ECM) via Filamin C ^29^. AMPK can also directly phosphorylate PDLIM5 at Ser177 to inhibit cell migration via Rho GEF 6 mediated suppression of Rac1 signalling ^30^. These pathways may therefore contribute to the dysregulated CREB signalling and upregulation of ECM proteins observed in C18ORF25 KO skeletal muscles ^11^.

Interestingly, the protein with the greatest differential binding to C18ORF25 in response to AMPK activation and lost following mutation of S66/67A was GNAS. Ubiquitously expressed, GNAS associates G-protein coupled receptors with adenylyl cyclase and is required for cAMP-dependent PKA signalling ^31^. We have previously shown that C18ORF25 is required for PKA-dependent signal transduction following contraction-dependent AMPK activation ^11^. Hence, the observation of differential enrichment of GNAS in the current data provide important mechanistic insights into the possible link between C18ORF25 and PKA-dependent signalling. Previous work by Chen et al. ^32^ showed that a germline haploinsufficiency of GNAS in mice resulted in obesity, glucose intolerance and insulin resistance. They also developed a skeletal muscle-specific G(s)alpha-knockout (KO) mice (MGsKO) which suffered from whole body glucose intolerance with no abnormalities in insulin secretion or sensitivity ^33^. Muscles from these mice also had reduced fatty acid oxidation despite a shift towards more oxidative fiber type distribution. Furthermore, *ex vivo* AICAR-stimulated glucose uptake was increased in MGsKO. Interestingly, there are several observed phenotypic similarities between MGsKO and C18ORF25-KO mice. For example, *soleus* muscles have reductions in tetanic force production, a reduction in lean mass and decreased muscle cross sectional area (CSA). Collectively, this suggests that healthy cAMP/PKA-dependent signalling in skeletal muscle may be mediated, at least partly, through C18ORF25 and its interactions with GNAS.

### Loss of C18ORF25 increases *ex vivo* contraction-stimulated glucose uptake

A large subset of proteins identified in the S67 phosphorylation-dependent interactome of C18OR25 comprised GTPases that have been implicated in GLUT4 trafficking. For example, Rab4 and Rab5 have been shown to localize to GLUT4 containing storage vesicles and are involved in their formation as well as shuttling to- and fusion with the plasma membrane ^34^. Rab5 has also been implicated in the insulin induced association of IRS1 with IR and P13K ^35^. The small GTPase Rac1 was also identified in the phosphorylation-dependent interactome and is required for GLUT4 translocation in skeletal muscle ^36^. In addition, Ras associated protein RALA was also identified which is interesting given that it localizes to GLUT4 vesicles and is required for insulin-stimulated GLUT4 trafficking ^37^. Furthermore, activation of β1 adrenergic receptors has been shown to activate Ras via CNrasGEF in a GNAS-dependent manner ^38^, and loss of GNAS in skeletal muscle increases contraction-stimulated glucose uptake ^33^. Given the observed phenotypic similarities between loss of GNAS and C18ORF25 (e.g reduced lean mass, CSA and muscle function), we next tested the role of C18ORF25 in contraction-stimulated glucose uptake. We harvested left and right EDL muscles from WT vs KO mice where the left muscle was kept in a basal state while the right muscle was subject to electrically stimulated contraction followed by metabolic tracing of radio-labelled ^3^H-2-deoxyglucose (2DOG) of both muscles. We observed elevated basal and contraction-stimulated 2DOG uptake (∼1.5x) in both male and female *C18Orf25* KO mice (**Figure 2A-B**). The contraction protocol increased the activity of AMPK to similar levels in both WT and KO mice as assessed by phosphorylation of S79 on ACC and T172 on AMPK (**Figure 2C**). It is also important to note that our previous proteomic analysis of skeletal muscle from WT and KO mice did not show any differences in the abundance of GLUT4 which might account for these differences in glucose uptake ^11^. While it is tempting to speculate that reduced PKA activity in the absence of C18ORF25 (as we have previously shown in ^11^ may be responsible for this phenotype, the relationship between cAMP/PKA signalling and skeletal muscle glucose uptake remains complex. Ortmeyer ^39^ highlighted the inhibition of PKA activity required for healthy insulin signalling in *vastus lateralis* muscles of rhesus monkeys while Sato et al. ^40^ reported that increased cAMP and therefore PKA activity promotes translocation of GLUT4 to the surface membrane via mTORC2 phosphorylation independent of Akt or AS160 in rat *soleus* muscles. Furthermore, Ngala et al. ^41^ demonstrate that different activators of β2-adrenoreceptors can have opposing effects on skeletal muscle cAMP levels and glucose uptake, attributing this divergence to ‘ligand-directed signalling’. Additional experiments are therefore required to elucidate the exact mode of action of C18ORF25 in the context of skeletal muscle glucose uptake.

**Figure 2.**
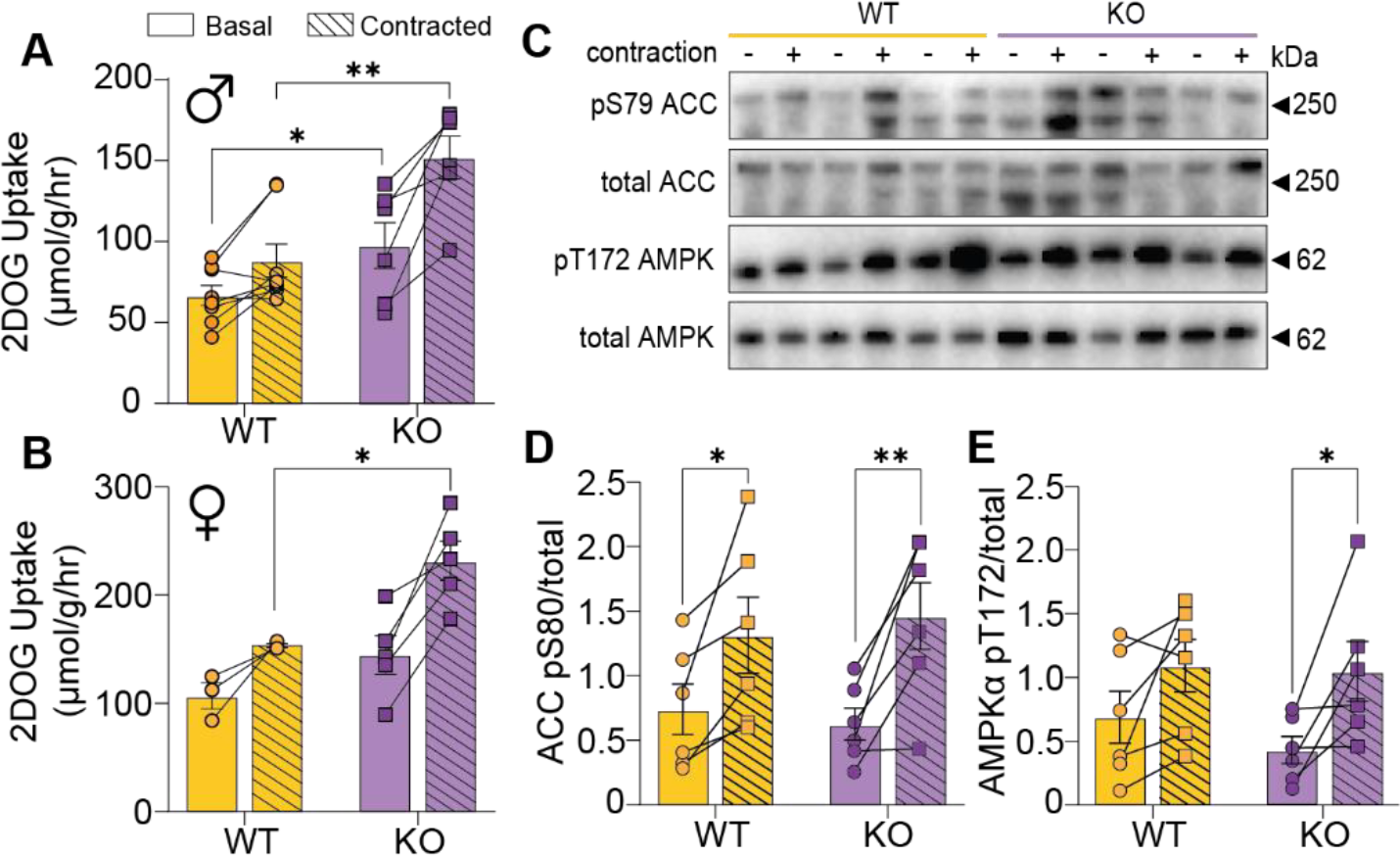
Contraction-stimulated 2-deoxyglucose (2DOG) uptake in *extensor digitorum longus* (EDL) muscles from wild type (WT) and *C18Orf25* knockout (KO) mice. 2DOG uptake in (**A**) males (n=6-7) and (**B**) females (n=3-5). (**C**) Representative image of western-blot analysis (n=3), and (**D-E**) densitometry quantification (n=6). *Student’s t-test; *p<0.05, **p<0.01

### C18ORF25 localises to the nucleus and sarcomere

Proteins interacting with C18ORF25 display an enrichment in localisation to the nucleus (e.g JUN, FOXK2 transcription factors), cytoskeleton (e.g RHOA, CDC42, Rac1), sarcolemma (e.g various integrins), and the sarcomere including Z-band (e.g FHL2 and PDLIM5) and A-band (e.g MYL2) (**Table S1**). To determine C18ORF25 localisation in skeletal muscle tissue, we next performed immunofluorescence microscopy. We injected *tibialis anterior* muscles with either AAV6:empty vector (EV) into the left leg or AAV6:N-FLAG-C18ORF25 into the right leg. Longitudinal sectioning and immunofluorescence microscopy revealed co-localisation to nuclear DAPI and interdigitation with Alpha-actinin 2, a Z-band enriched protein involved in cross-linking or sarcomeric actin filaments (**Figure 3A**). We also observed localisation to the sarcolemma consistent with the association to many plasma membrane proteins. Taken together, these data are highly consistent with the subcellular localisation of proteins identified in AP-MS and further suggest broad functions of C18ORF25.

**Figure 3A.**
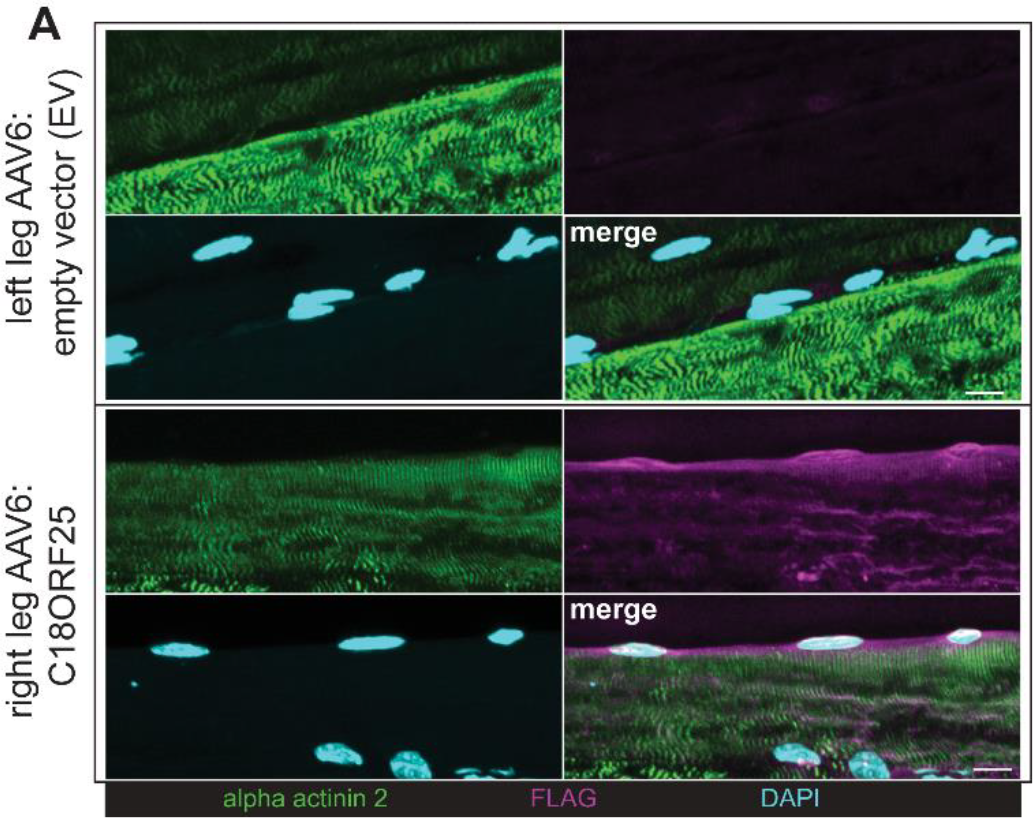
Subcellular localisation analysis via immunofluorescence microscopy of *tibialis anterior* muscles expressing empty vector or N-FLAG-C18ORF25 into the right leg. Scale bar = [10 µm]

### Integrated single fiber physiology and single fiber proteomics

We have previously shown that loss of C18ORF25 results in fiber-type specific defects in calcium handling of *soleus* muscle ^11^. To further investigate these fiber-type specific mechanisms, we developed an integrated single fiber physiology and single fiber proteomic pipeline (**Figure 4A**). Firstly, single muscle fibers were isolated and skinned by mechanically removing the sarcolemma. Next, the skinned single fiber was attached to a micro-force transducer, and various steps in the excitation-contraction (EC) pathway were assessed including assessment of Ca^2+^-sensitivity of the contractile apparatus, the maximum Ca^2+^-activated force response, and evaluation of sarcoplasmic reticulum (SR) Ca^2+^-loading, release, and leak properties. Finally, each single fiber was classified as either fast-twitch (FT) or slow-twitch (ST) by exposure to strontium and then snap frozen for single fiber proteomic analysis. A unique feature of this pipeline is the ability to measure phenotypic responses of the EC pathway and perform proteomic quantification in the exact same cell which provides opportunities to identify phenotype:protein associations. It’s important to note that; i) fibers skinned via mechanical means retain functional intracellular components necessary for normal EC coupling, and (ii) because fibers are skinned under paraffin oil, the SR remains intact and the calcium handling properties can be assessed ^16^.

**Figure 4.**
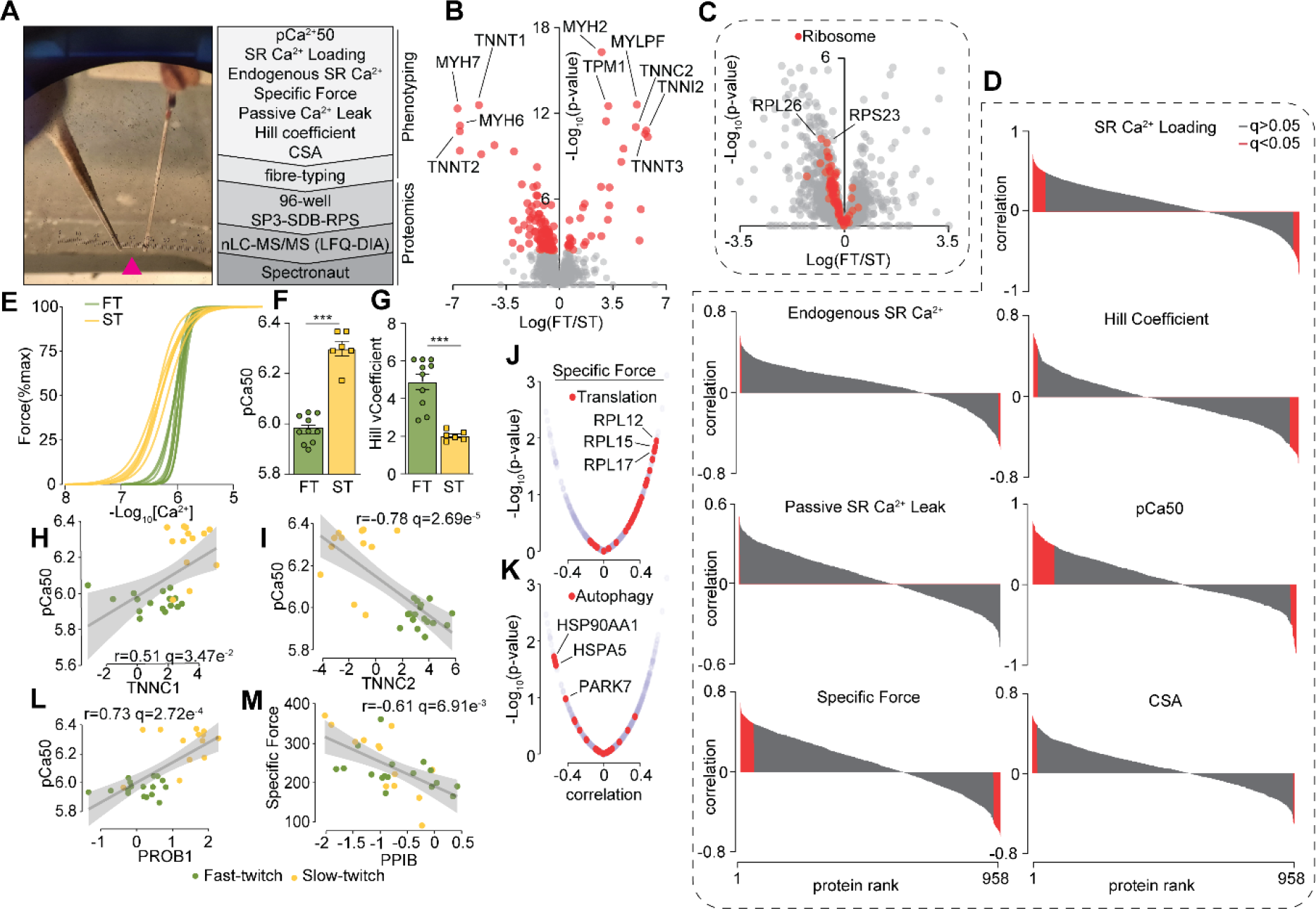
Integrated single fiber physiology and single fiber proteomic platform. (**A**) Photo of a single mechanically-skinned muscle fiber (pink arrow) sutured onto the force transducer and clamped between forceps and overview of the phenotypic assessments and proteomic workflow. (**B**) Volcano plot highlighting proteins quantified in fast-twitch (FT) vs slow-twitch (ST) muscle fibers. (**C**) Quantification of ribosomal proteins in FT vs ST muscle fibers. (**D**) Correlation of protein abundance to SR Ca2+ Loading, endogenous SR Ca2+, hill coefficient, passive SR Ca2+ leak, pCa50, specific force and CSA. (**E**) Contractile force of individual FT and ST muscle fibers during calcium dose response, and (**F**) calculated pCa50, and (**G**) Hill Coefficient. Correlation of (**H**) TNNC1, and (**I**) TNNC2 protein abundance to pCa50. Correlation of proteins annotated as (**J**) “Translation” and (**K**) “Autophagy” in KEGG/Reactome to specific force. (**L**) Correlation of PROB1 protein abundance to pCa50, and (**M**) correlation of PPIB protein abundance to specific force.

The integrated single fiber physiology and paired proteomic pipeline was applied to the analysis of 32 individual fibers from WT and KO *soleus* muscles resulting in the quantification of 1,612 proteins (**Table S2**). We first focused our analysis of proteins significantly regulated between FT and ST fibers independent of genotype. A total of 138 proteins were significantly regulated between these two fiber-types (**Figure 4B**). All of these have been identified in previous single fiber proteomic analyses and include upregulation of contractile protein isoforms in FT fibers such as MYH2, MYLPF, TNNI2, TNNT3, MYOM2 and MYL1, and proteins upregulated in ST fibers such as TNNT1, MYH6, MYH7 and TNNT2 ^42^; ^43^; ^44^; ^45^. Gene set enrichment analysis (GSEA) identified an enrichment of mitochondrial processes such as TCA cycle and mitochondrial translation in ST fibers while pathways related to glycogenolysis were enriched in FT fibers which is consistent with previous studies described above (**Table S3**). We also observed a general enrichment of ribosomal proteins in ST fibers with a ∼50% increase in RPS15A, RPS23, RPL26, RPL28, RPS29 and RPL35 which have recently been described in fiber heterogeneity (https://www.biorxiv.org/content/10.1101/2023.09.07.556665v1) (**Figure 4C**).

Next, we correlated each of the phenotypes to each quantified protein in a pairwise manner. After filtering (see methods), a total of 752 significant protein:phenotype associations were identified (**Figure 4D**)(**Table S4**). Classically, ST type-I fibers have slower shortening velocity while FT type-IIa/x/b fibers generate higher velocities. These contractile properties are a result of several phenotypic differences including the sensitivity of the sarcomere to Ca^2+^, differences in calcium loading and unloading into the SR, structural changes, and metabolic differences, much of which arise due to differential expression of contractile protein isoforms specific to each fibre type ^46, 47, 48^. For example, ST fibers have increased sensitivity to Ca^2+^ as calculated by the concentration of Ca^2+^ needed to elicit 50% of maximal force, or pCa50, and a lower Hill Coefficient (**Figure 4F-G**) ^47^. As such, we observed significant positive correlation of the slow troponin C isoform (TNNC1), and a significant negative correlation of the fast troponin C (TNNC2) with pCa50 (**Figure 4H-I**). Although the different troponin isoforms have well characterised sensitivities to Ca^2+^, it is important to note that the observed protein:phenotype correlations listed in **Table S4** may simply arise due to fiber-type phenotypic differences and may not be causal in nature. For example, the slow twitch myosin isoform MYH7, which does not confer Ca^2+^ sensitivity, follows a similar correlation profile with TNNC1 with respect to pCa50. Despite this limitation, we observed several other fascinating observations. For example, a GSEA analysis was performed on the protein associations to specific force which identified positively and negatively correlated to proteins involved in translation and autophagy, respectively (**Figure 4J-K**). Fibers with high specific force had elevated levels of ribosomal proteins while fibers with low specific force had elevated levels of various proteins involved in proteostasis including chaperones and PARK7. We also observed additional novel associations such as a positive correlation between PROB1 and pCa50 (**Figure 4L)**. PROB1 (also known as C5ORF65) is a proline-rich nucleocytoplasmic protein with unknown function. Genetic variants in human *PROB1* have been implicated in keratoconus, an eye disease resulting from thinning of the cornea ^49^. Its role in the EC pathway is unknown but its expression is highly enriched in striated muscle which warrants further investigation (https://www.proteinatlas.org/ENSG00000228672-PROB1/tissue). Another interesting association is the negative correlation of PPIB with specific force (**Figure 4M**). PPIB (also known as Cyclophilin B; CYPB) is an SR-localised chaperone and member of the family of Peptidyl-prolyl cis-trans isomerases. The protein has been primarily studied in the context of procollagen modification, sorting and transport through the secretory compartment. Mice lacking *Ppib* have several musculoskeletal defects and genetic variants in human *PPIB* cause osteogenesis imperfecta, a connective tissue disease characterized by developmental defects and bone fragility ^50^. The observed association between PPIB and contractile function within these single fiber experiments warrants further investigation, especially since it associates with other proteins in addition to collagens such as FHOD1, a Rho-ROCK-dependent factor required for F-actin assembly ^51^.

### Fiber-type specific differences in *C18Orf25* knockout mice

Loss of C18ORF25 results in a decrease in the amount of Ca^2+^ that can be loaded into and released from the SR in fast-twitch (FT) but not slow-twitch (ST) muscle fibers (**Figure 5A**). Furthermore, the levels of passive Ca^2+^ leak from the SR is significantly elevated in FT but not ST fibers of KO mice (**Figure 5B**). It is important to note that loss of C18ORF25 does not result in changes in the abundance or the force-generating properties of the contractile apparatus, and there is no difference in the distribution of fiber-types as assessed by immunofluorescence microscopy ^11^. We next investigated the fiber-type specific differences in calcium handling by comparing the FT and ST proteomes of WT vs KO *soleus* muscles. A total of 36 proteins were differentially regulated between WT and KO FT fibers (q<0.05) while no proteins were regulated in ST fibers, consistent with no phenotypic differences in the latter (**Figure 5C-D**)(**Table S2**). The most significantly up-regulated protein was COBL, a Ca^2+^/calmodulin-dependent actin nucleator. COBL was originally shown to be enriched in the central nervous system (CNS) ^52^ but has more recently been established as a striated-muscle enriched protein (https://www.proteinatlas.org/ENSG00000106078-COBL/tissue). Similar to C18ORF25, COBL is also localised to the Z-band ^53^ but was not identified in the interactome via AP-MS. COBL shows a trend for positive correlation to Ca^2+^ SR loading but is not significant when adjusting for multiple hypothesis testing (**Figure 5E**)(**Table S4**). The protein with the largest significant fold-change was OPLAH, a 5-oxoprolinase that is enriched in the testes, gall bladder and striated muscle tissue (https://www.proteinatlas.org/ENSG00000178814-OPLAH/tissue). It is unclear how OPLAH might be associated with the observed phenotypic differences in FT fiber calcium handling, and it was not identified in enough replicates for protein:phenotype correlation analysis. However, it is interesting to note that some patients with OPLAH deficiency have calcium oxalate/carbonate urolithiasis ^54^. Another interesting protein down-regulated in FT fibers of *C18Orf25* KO mice is AHNAK, a ubiquitously expressed chaperone protein involved in regulating Ca^2+^ release via L-type voltage gated Ca^2+^ channels (LVGCC) ^55^. In cardiomyocytes, AHNAK has been shown to be phosphorylated by PKA which results in its dissociation from LVGCCs, therefore relieving inhibition of the channel and promoting calcium flux ^56^. It is thought to play a similar role in skeletal muscle excitation-contraction coupling (ECC) where AHNAK has been shown to colocalize with t-tubular LVGCCs ^57^. One of the most interesting proteins significantly different between WT- and KO-FT fibers was CAMK2G which was also positively correlated with SR Ca^2+^ loading (**Figure 5F**). Indeed, CAMK2 inhibition in fast-twitch fibers inhibits Ca^2+^ release ^58^ and is known to phosphorylate several proteins localised to the SR ^59^. Taken together, our integrated single fiber phenotypic and single fiber proteomic analysis has provided vital insights into the fiber-type specific regulation of calcium handling following loss of *C18Orf25*.

**Figure 5.**
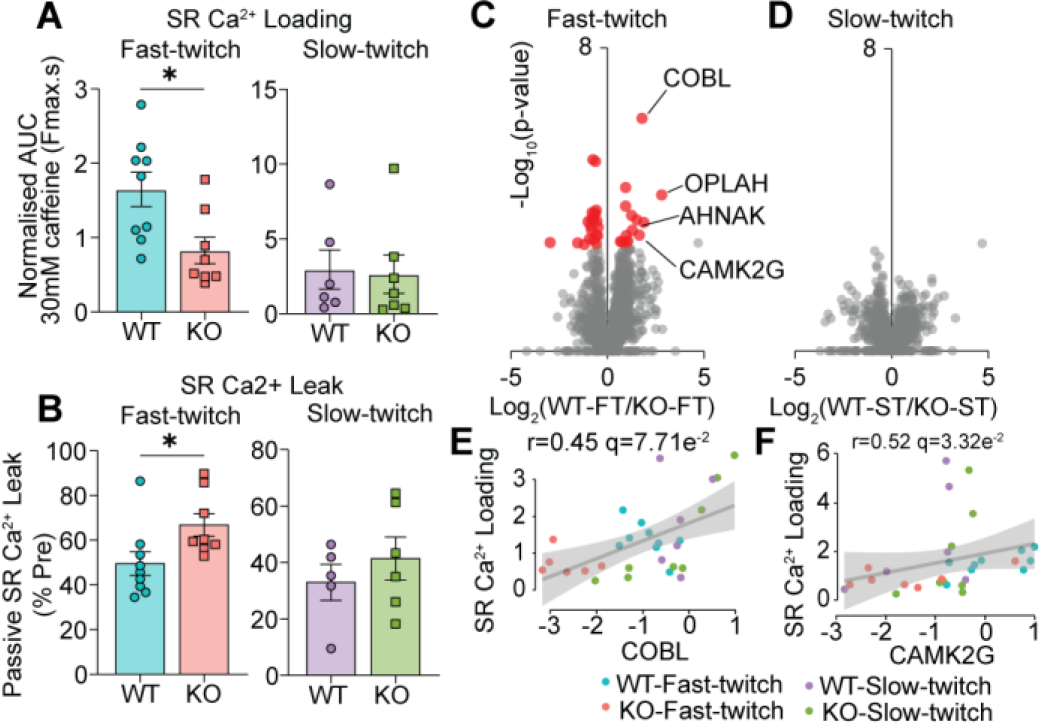
Phenotyping and proteomic profiling of single fibers from *soleus* muscles of wildtype (WT) vs C18Orf25 knockout (KO). (**A**)Sarcoplasmic reticulum (SR) calcium (Ca^2+^) loading ability and (**B**) passive SR Ca^2+^ leak of fast twitch (FT) and slow twitch (ST) fibers from wildtype (WT) vs knockout (KO) muscles. (**C**) Volcano plot highlighting proteins quantified in FT fibers and (**D**) ST fibers from WT vs KO muscles. (**E**) Correlation of COBL protein abundance and (**F**) CAMK2G protein abundance to SR Ca^2+^ loading.

## Conclusion

We were able to identify interacting partners of C18ORF25 via AP-MS. GNAS, a subunit of GPCRs, appeared as an exciting interacting partner significantly enriched in response to AMPK activation and attenuated following mutation of S66/67A. It is tempting to suggest that C18ORF25 may exert its function via interaction with GNAS, and that the absence of C18ORF25 therefore results in impaired PKA dependent signalling via disrupted coupling of adenylyl cyclase with GPCRs. Several candidates of interest identified in our interactomics and single fiber proteomics data further suggest a major involvement of C18ORF25 in SR Ca^2+^ handling and GLUT4 trafficking, which is reflected in our *ex vivo* contractile studies and exercise-induced glucose uptake experiments. Another highlight of this study was the ability to correlate over 300 proteins with single fiber phenotypes such as force output, Ca^2+^ handling and Ca^2+^ sensitivity of the contractile apparatus. Here, most proteins differentially regulated in WT vs KO fast-twitch fibers were also found to correlate with these parameters. Additionally, immunofluorescent staining revealed C18ORF25 localized to the Z-band and nuclei. Overall, we were able to utilize various methods of molecular analyses to reveal further direct and indirect mechanistic insights of C18ORF25.

